# Extracellular vesicle formation in *Cryptococcus deuterogattii* impacts fungal virulence and requires the *NOP16* gene

**DOI:** 10.1101/2022.06.06.494995

**Authors:** Rafael F. Castelli, Alana Pereira, Leandro Honorato, Alessandro Valdez, Haroldo C. de Oliveira, Jaqueline M. Bazioli, Ane W. A. Garcia, Tabata D’Maiella Freitas Klimeck, Flavia C. G. Reis, Charley C. Staats, Leonardo Nimrichter, Taícia P. Fill, Marcio L. Rodrigues

## Abstract

Small molecules are components of fungal extracellular vesicles (EVs), but their biological roles are only superficially known. *NOP16* is a eukaryotic gene that is required for the activity of benzimidazoles against *Cryptococcus deuterogattii*. In this study, during the phenotypic characterization of *C. deuterogattii* mutants lacking *NOP16* expression, we observed that this gene was required for EV production. Analysis of the small molecule composition of EVs produced by wild-type cells and two independent *nop16*Δ mutants revealed that the deletion of *NOP16* resulted not only in a reduced number of EVs but also an altered small molecule composition. In a *Galleria mellonella* model of infection, the *nop16*Δ mutants were hypovirulent. The hypovirulent phenotype was reverted when EVs produced by wild-type cells, but not mutant EVs, were co-injected with the *nop16*Δ cells in *G. mellonella*. These results reveal a role for *NOP16* in EV biogenesis and cargo, and also indicate that the composition of EVs is determinant for cryptococcal virulence.

## Introduction

Extracellular vesicles (EVs) are membranous structures produced by all domains of life (1). In microorganisms, EVs participate in processes of immunopathogenesis, cell to cell communication, cellular differentiation, and antimicrobial resistance, among others [reviewed in (2)]. Fungal EVs were first characterized in the *Cryptococcus* genus (3). In this model, EVs have immunostimulatory activity (4) and vaccinal potential (5). Cryptococcal EVs are also required for fungal virulence (6) and intraspecies communication (7).

As widely discussed in the literature, several questions related to fungal EVs remain unknown (8). The mechanisms of biogenesis of fungal EVs are still poorly known, which impairs the design of experimental models for the study of their functions. Although much progress has been made in the identification of the protein and nucleic acid components of fungal EVs (5, 9–12), their small molecule composition was only recently addressed in *C. deuterogattii* (13), *Penicillium digitatum* (14), *P. chrysogenum* (15), and *Histoplasma capsulatum* (16). In *C. deuterogattii*, small molecule analysis of EVs revealed the presence of a peptide controlling infection in the *Galleria mellonella* model (13). A possible relationship between small molecule composition and fungal virulence remains to be determined, as well as the genes regulating the formation of fungal EVs and their small molecule cargo.

*NOP16* is the gene encoding nucleolar protein 16. In humans, Nop16 is expressed in 237 different tissues (17), but its functions are poorly known. In *Saccharomyces cerevisiae*, Nop16 is a constituent of 66S pre-ribosomal particles, with involvement in 60S ribosomal subunit biogenesis (18, 19). There is no evidence in the literature pointing to the roles of Nop16 in the formation of EVs, although ribosomal proteins are abundant in cryptococcal EVs (5, 20).

During the search for novel anti-cryptococcal agents, we found that Nop16 was involved in the antifungal activity of mebendazole by still unknown mechanisms (21). In this study, we report on the generation and characterization of two independent mutants of *C. deuterogattii* lacking expression of Nop16. We found no connections between antifungal activity and Nop16 expression. However, in comparison to wild-type (WT) cells, the *nop1*6Δ mutants were hypovirulent in a *G. mellonella* model of infection, although they shared many phenotypic properties with WT cells including growth rates, ultrastructural features, capsule formation, and susceptibility to phagocytosis. The *nop1*6Δ mutants were less efficient in producing EVs, in association with an altered small molecule cargo. Infection of *G. mellonella* with the *nop1*6Δ mutant strains in the presence of EVs produced by parental cells resulted in increased virulence, suggesting a role for *NOP16* in EV biogenesis and cargo. These results also suggest that the composition of EVs can be determinant for the virulence of *C. deuterogattii*.

## Results

### Generation of the *nop1*6Δ mutants

We generated null mutants of the *NOP16* (CNBG_3695) gene in *C. deuterogattii* by employing the Deslgate methodology, as previously described (22). Two null mutants were identified based on the absence of the amplification with diagnostic primers (Fig 1A and 1B) and decreased sensitivity to mebendazole (Fig. 1C), based on the previously described association between the antifungal activity of benzimidazoles and *NOP16* expression (21).

**Figure 1:**
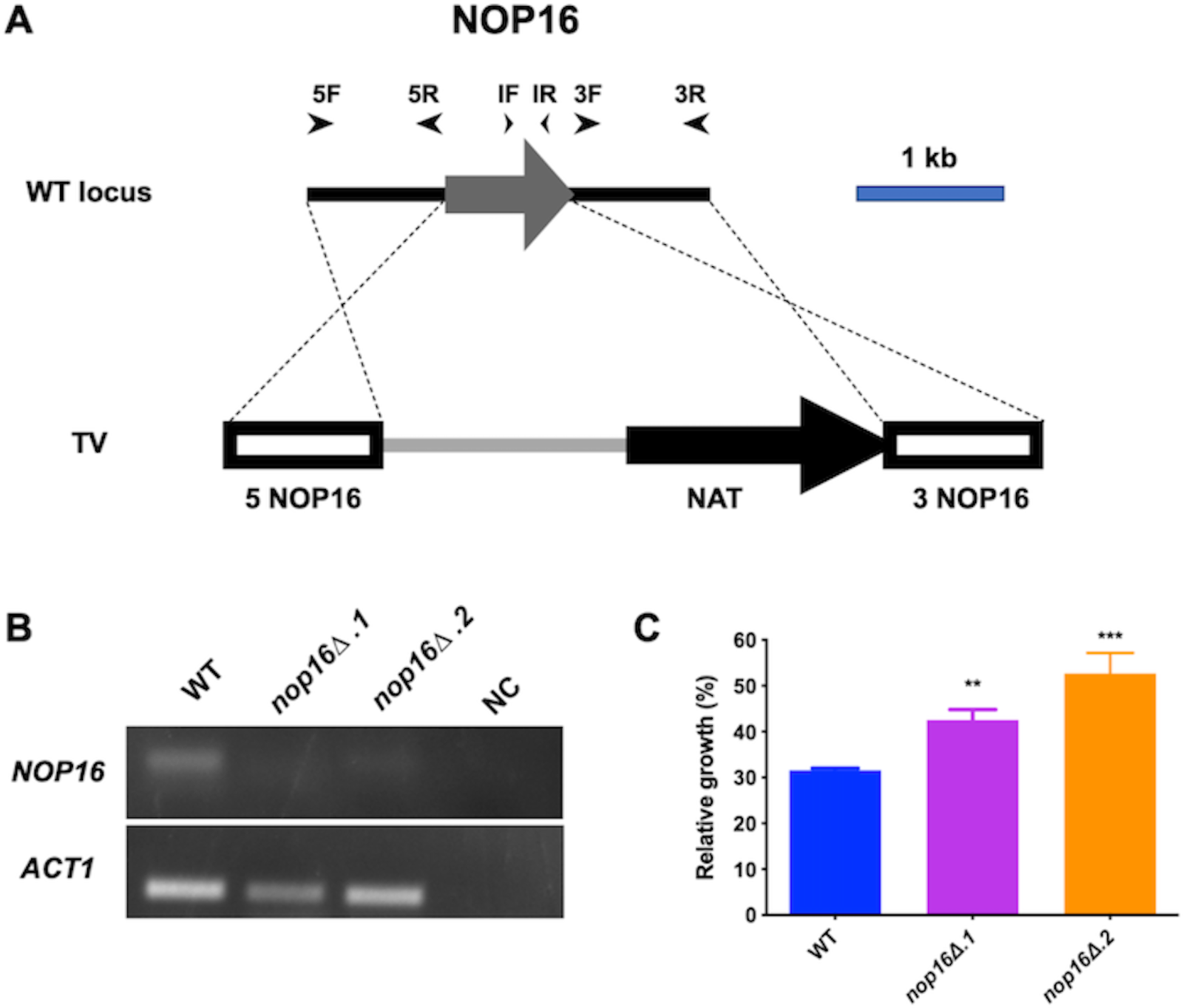
Generation of null mutants of *NOP16* in *C. deuterogattii*. (A) *NOP16* inactivation strategy. TV is the representation of the targeting vector constructed by the Delsgate methodology, with 5 NOP16 and 3 NOP16 representing the 5’ and 3’ gene flanks of the *NOP16* gene, respectively. The primers used to amplify the 5’ (5F and 5R) and 3’ (3F and 3R) of *NOP16* are represented as arrowheads. NATR is the cassette that confers noursethricin resistance. (B) Nurseothricin-resistant cells were evaluated for the presence of the *NOP16* gene using internal diagnostic primers (IF and IR), using the *ACT* gene as a loading control. (C) WT and two null mutants were evaluated for their sensitivity to mebendazole. Bars represent the average of the ratio between growth in 1 mM mebendazole normalized to the growth in a drug-free medium obtained in three independent experiments. Mutant cells display decreased sensitivity to mebendazole (** *P < 0.01 and *** P < 0.0001)* as revealed by ANOVA followed by Dunnett multiple comparison analysis.

### Phenotypic characterization of *nop1*6Δ.1 and *nop1*6Δ.2 mutant strains revealed a hypovirulent profile

As part of our antifungal development program (21), we tested possible links between Nop16 and antifungal activity but did not find any clear connection (data not shown). Nevertheless, we proceeded with the characterization of the mutants. Our tests included virulence potential in the *G. mellonella* model, determination of growth rates, observation of general cellular aspects, production of virulence factors, and characterization of EVs, as follows below.

We first tested the virulence potential of the mutants by comparing their ability to kill *G. mellonella* with that demonstrated by parental (wild-type, WT) cells. Both mutants took longer than WT cells to kill the invertebrate host (Figure 2, P < 0.0001). In the representative experiment (1 out of 3) that we describe in this study, all larvae were killed by WT cells on day 2 post-infection, while the mutants took 7 days to kill the whole *G. mellonella* population. Replicates produced similar results (data not shown).

**Figure 2.**
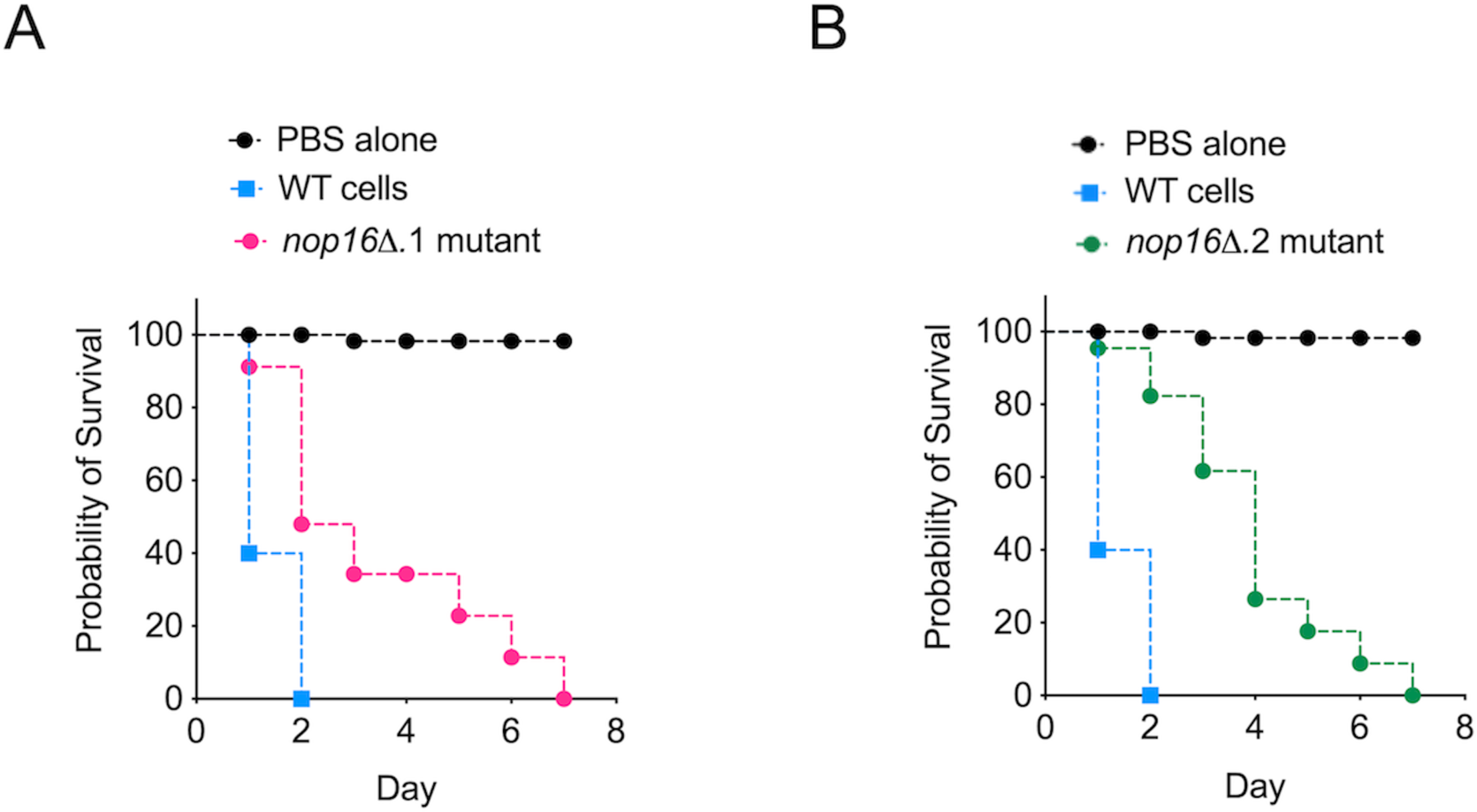
Deletion of Nop16 led to a hypovirulent phenotype in *C. deuterogattii*. Infection of *G. mellonella* with the independent *nop1*6Δ mutants 1 (A) and 2 (B) resulted in a smaller efficacy in killing the animals, in comparison with wild-type (WT) cells (P < 0.0001 for both mutants). Control systems were injected with PBS only. Statistical analysis was performed with the Mantel-Cox test.

We asked why the *nop1*6Δ.1 and *nop1*6Δ.2 mutant strains had reduced abilities to kill *G. mellonella*. To address this question, we first determined the proliferation rates of the *nop1*6Δ.1 and *nop1*6Δ.2 mutants and compared them to those obtained for WT cells at 37°C in three different media. In all media (Sabouraud, YPD, and RPMI, Figure 3A-C), the mutants manifested growth rates that were higher than those of WT cells, which led us to discard the possibility that their decreased virulence was a consequence of reduced growth capacities. We then evaluated whether *NOP16* deletion induced evident cellular alterations in *C. deuterogattii*. WT cells and the *nop1*6Δ.1 and *nop1*6Δ.2 mutant strains had similar ultrastructural aspects, as concluded from the observation of transmission electron microscopy (TEM) micrographs (Figure 3D-F).

**Figure 3.**
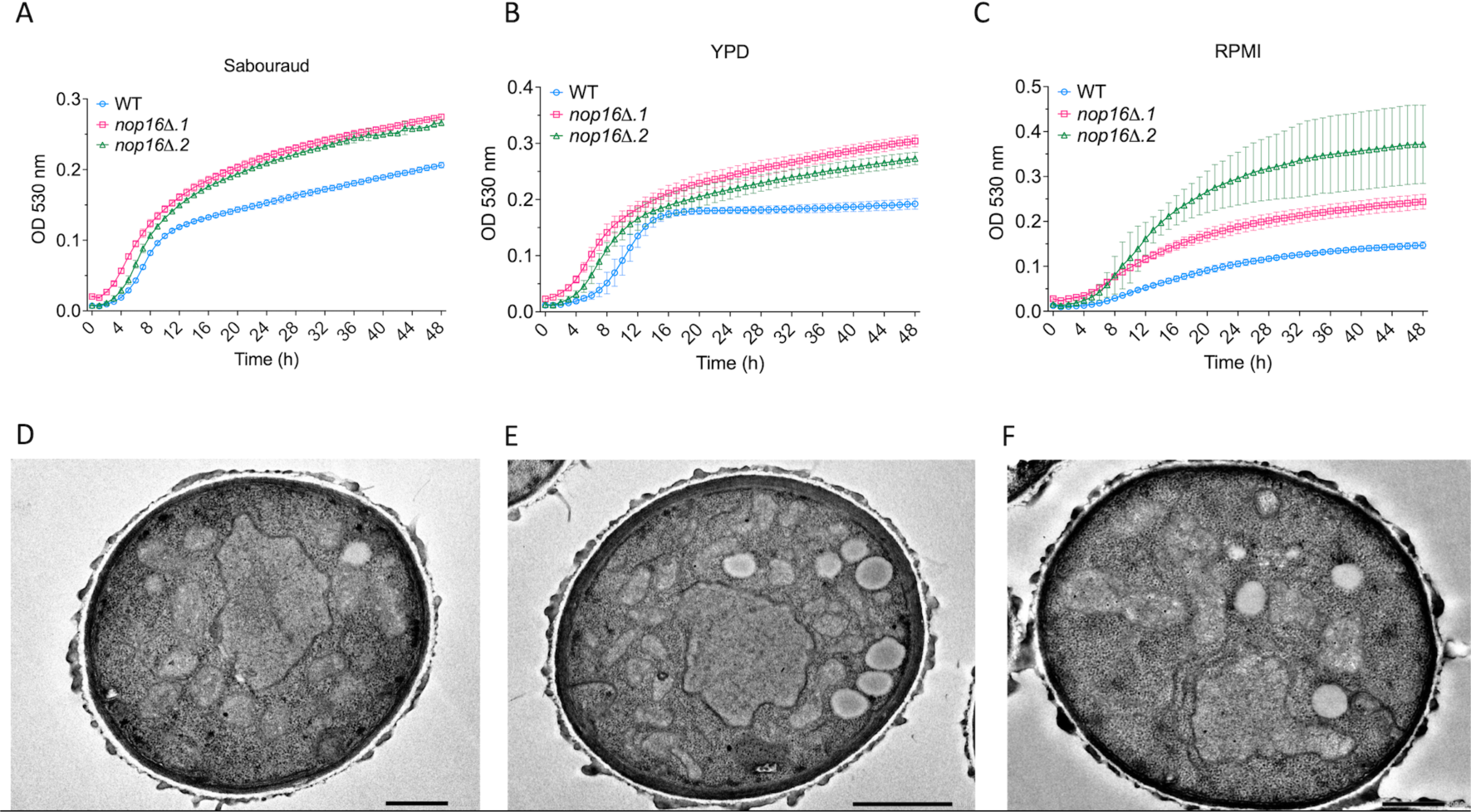
Analysis of growth rates (A-C) and ultrastructural aspects (D-F) of wild type (WT) and mutant cells lacking Nop16 (*nop1*6Δ.1 and *nop1*6Δ.2 strains). Independently on the use of Sabouraud (A), YPD (B), or RPMI (C) as the growth media, the mutant strains always manifested higher proliferation rates, in comparison to WT cells. Analysis of the ultrastructural features of WT (D), *nop1*6Δ.1 (E), and *nop1*6Δ.2 (F) cells revealed that Nop16 deletion resulted in no evident alterations. Scale bars in D-F correspond to 500 nm.

Since the cell surface of *C. deuterogattii* is essential for interaction with host immune cells and, consequently, pathogenesis (23), we evaluated whether *NOP16* deletion affected cell wall and capsular structures. Staining of the cell wall components chitin and chitooligomers of WT and mutant strains revealed similar characteristics (Figure 4A-C), which were in agreement with those reported in the literature (24). As for the capsular structures of *C. deuterogattii*, immunofluorescence analysis of parental and *nop1*6Δ.1 and *nop1*6Δ.2 cells using a monoclonal antibody to the main cryptococcal capsular component, namely glucuronoxylomannan (GXM), showed similar profiles of serological reactivity in all strains (Figure 4A-C). The mutants tended to produce higher amounts of extracellular GXM, but the differences were very discrete (Figure 4D). The general similarity in the capsular structures was confirmed by India ink counterstaining and no visual evidence of altered capsules was observed (Figure 4E-G). We then determined the capsular dimensions in the 3 strains. Although the capsules of the mutants were statistically smaller than those of WT cells (P < 0.0001), the average values were very close (Figure 4H). Scanning electron microscopy confirmed the immunofluorescence and counterstaining results, with no apparent differences between the capsules of WT and mutant cells (Figure 4I-K). Since the susceptibility to phagocytosis, which also involves the participation of the cryptococcal capsule (25), is a determinant for the pathogenesis of *Cryptococcus* (26), we also compared the phagocytic rates of WT and mutant strains. We found no significant differences between the phagocytic rates of WT and mutant cells by mouse macrophages (Figure 4L). Altogether, these results led us to the conclusion that the hypovirulent phenotypes of the *nop1*6Δ.1 and *nop1*6Δ.2 mutant strains were not related to evident cellular alterations, capsule formation nor its possible interference with phagocytosis. We also tested melanin formation as another cryptococcal virulence factor (27). Once again, WT and mutant strains had similar abilities to make pigments (data not shown).

**Figure 4.**
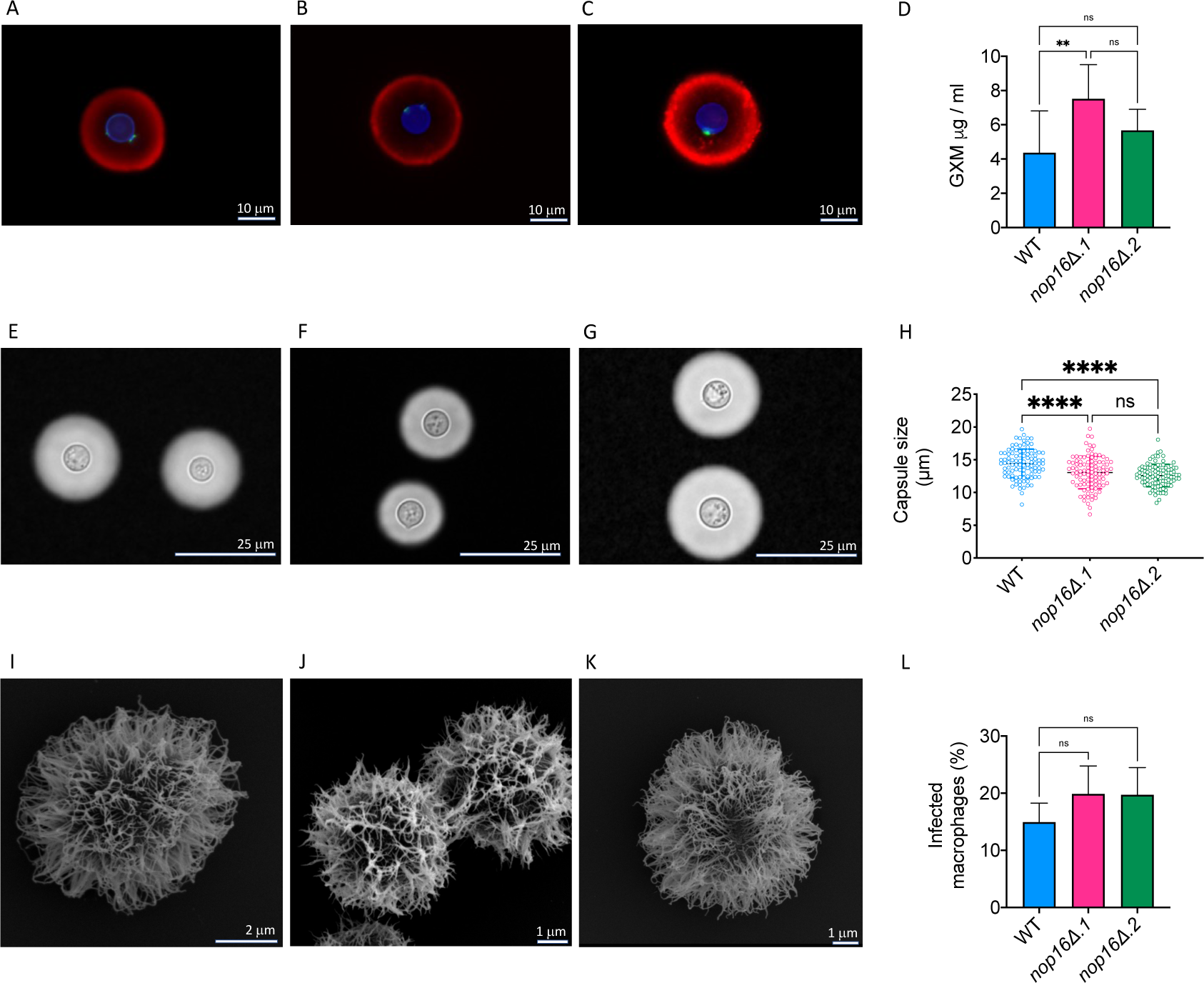
Analysis of the role of Nop16 in the surface architecture of *C. deuterogattii* and its impact on fungal phagocytosis. Fluorescence microscopy analysis of the cell surface of WT (A), *nop1*6Δ.1 (B), and *nop1*6Δ.2 (C) cells revealed similar aspects of cell wall chitin (blue fluorescence), chitooligomers (green fluorescence), and the capsule (red fluorescence) in the 3 strains. GXM secretion tended to be higher in the *nop1*6Δ.1 mutant (**P = 0.0053), but not the *nop1*6Δ.2 strain (D; ns, not significant). India ink counterstaining of WT (E), *nop1*6Δ.1 (F), and *nop1*6Δ.2 (G) cells suggested similar capsular dimensions, but the determination of the capsule sizes revealed lower average values for the mutant cells (H, ****P < 0.0001, with a 95% confidence level of 95.61%). SEM was also used for the observation of the capsules of WT (I), *nop1*6Δ.1 (J), and *nop1*6Δ.2 (K) cells, revealing similar capsular structures. The phagocytosis rates of the three strains by mouse macrophages were also determined (L). No significant (ns) differences between the three strains were observed.

### EV formation is affected in the *nop1*6Δ.1 and *nop1*6Δ.2 mutant strains

EVs are required for virulence mechanisms in *C. deuterogattii* (6). We then hypothesized that EV formation could be affected by *NOP16* deletion. The genes regulating formation of fungal EVs are not well known, but in *Saccharomyces cerevisiae* EV production was influenced by the functionality of Golgi-related secretory proteins (28). Therefore, we first analyzed the Golgi of WT and mutant strains after staining the cells with C6-NBD-ceramide (29).

WT cells showed the typical pattern of disperse Golgi staining previously observed in *C. neoformans* (30, 31) (Figure 5A). In mutant cells, Golgi staining was clearly less intense, and most of the cells gave weak or negative fluorescent signals. This visual perception was confirmed by quantitative determination of fluorescence staining, which revealed a significantly reduced fluorescence signal in mutant cells (Figure 5B). These results were suggestive that *NOP16* deletion somehow resulted in defects in the Golgi. We then asked whether EV formation was also altered in *nop16Δ* cells.

**Figure 5.**
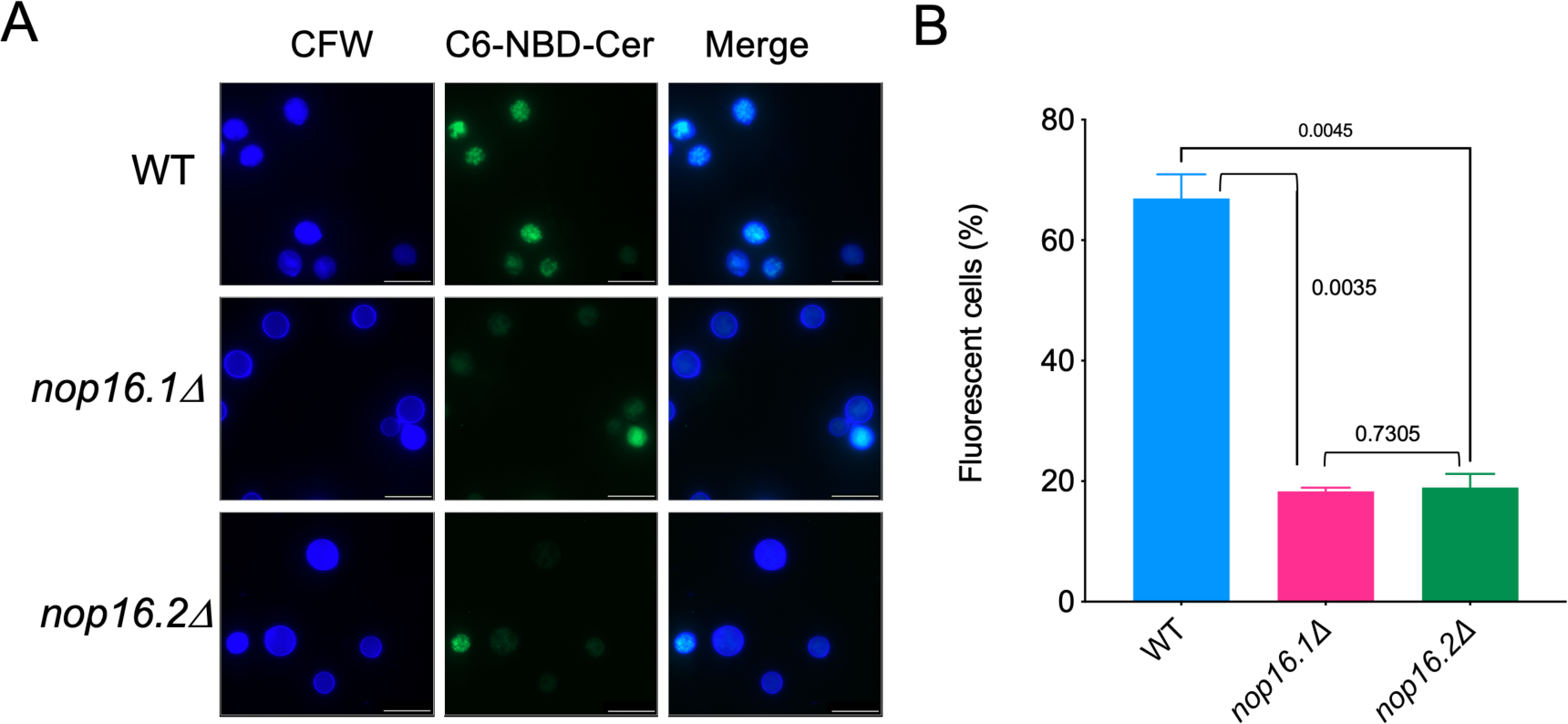
Cell wall (calcofluor white, CFW) and Golgi staining (C6-NBD-Cer) in WT and mutant cells. A. Microscopic examination of stained fungal cells suggested reduced levels of Golgi staining in mutant cells. B. Quantitative determination of fluorescent cells (100 cells for each condition) confirmed a significantly reduced detection of Golgi-staining in the mutants, in comparison with WT cells. Scale bars, 10 µm.

To address this question, we isolated EVs from WT and mutant cells for characterization by TEM and nanoparticle tracking analysis (NTA) (Figure 6A). TEM revealed that WT and mutant cells produced EVs with their typical morphological aspects, including cup-shaped structures and bilayered membranes. NTA demonstrated that in the 3 strains, the EV population was mainly distributed in the 100-300 nm range, as extensively observed for the *Cryptococcus* genus (5, 32, 33). Minor populations in the 300-600 nm range were also observed in the three strains. However, the quantification of EVs produced by the three strains by NTA revealed a significantly reduced formation of vesicles in the mutant strains (Figure 6B). After normalization of the number of EVs in each sample to the number of EV-producing cells, the *nop1*6Δ.1 mutant was approximately twice less efficient than WT cells to produce EVs, while EV production was approximately 3-fold higher in the parental strain than in the *nop1*6Δ.2 mutant. In an attempt to validate this observation, we measured extracellular urease activity in the 3 strains, since this enzyme is an extracellular virulence factor exported in EVs (20). Urease activity was significantly smaller in mutant cells, in comparison with the enzyme activity measured in the parental strain (Figure 6C).

**Figure 6.**
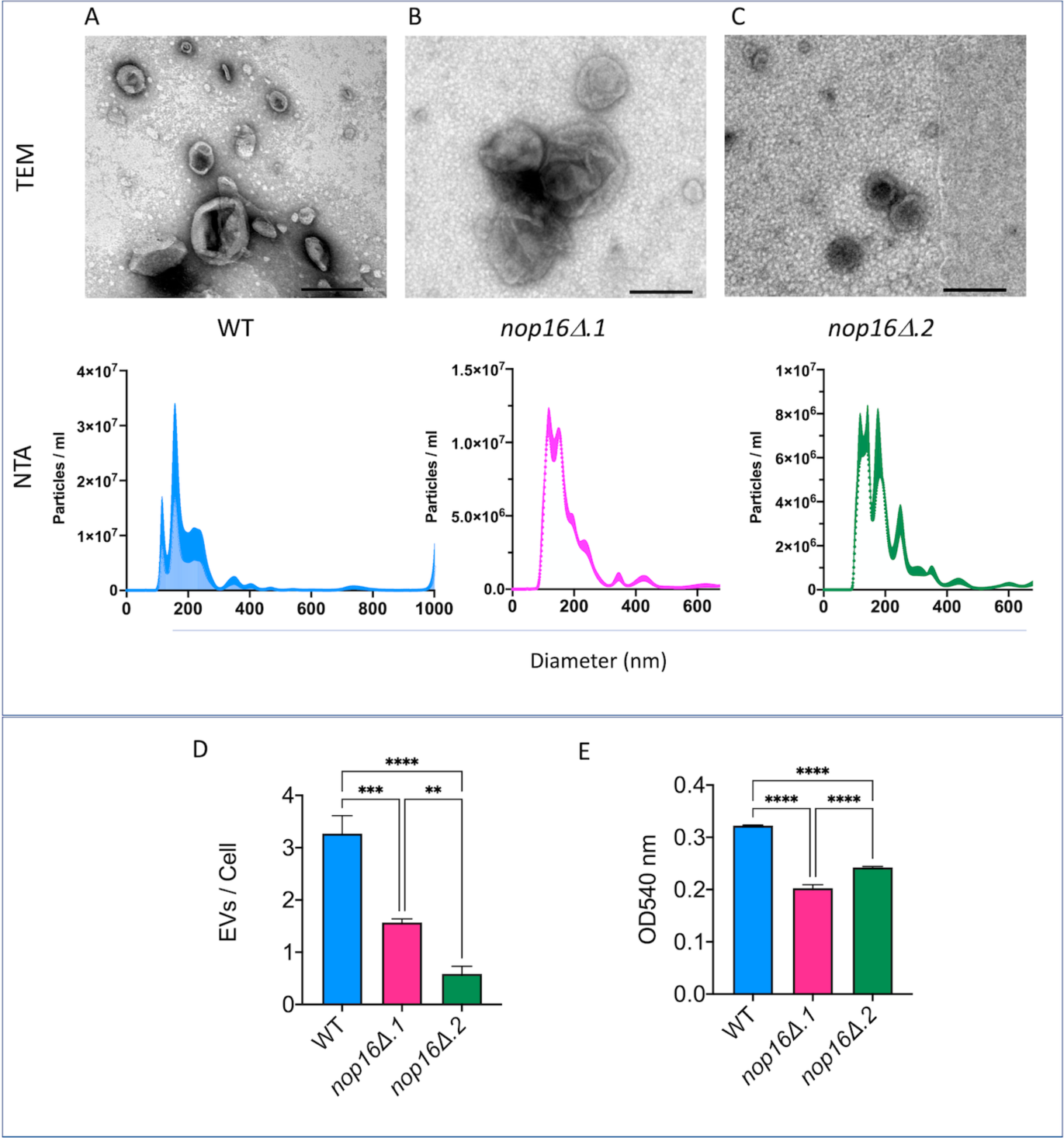
Analysis of the impact of Nop16 deletion on EV formation in *C. deuterogattii*. EVs produced by wild-type (A), *nop1*6Δ.1 (B) and *nop1*6Δ.2 (C) cells were characterized by TEM (upper panels) and NTA (lower panels). EVs from all strains, in general, manifested similar properties. D. Quantification of EVs produced by each strain revealed a significantly reduced number of vesicles produced by mutant cells, in comparison to the WT strain (****P < 0.0001; ***P = 0.002). EV production was smaller in the *nop1*6Δ.2 mutant than in the *nop1*6Δ.1 strain (**P = 0.0036). E. Measurement of urease activity in the 3 strains revealed that *NOP16* deletion resulted in a lower ability to hydrolyze urea (****P < 0.0001). Both mutants manifested similar urease activities (ns, not significant).

### *NOP16* deletion affects the small molecule composition of EVs

The reduced formation of EVs in the *nop1*6Δ.1 and *nop1*6Δ.2 mutant strains suggested a role for Nop16 in EV biogenesis. Since EV biogenesis involves the mechanisms of cargo (34), we also asked whether the composition of EVs was altered after *NOP16* deletion. As a proof of concept, we focused our analysis on the 13 EV small molecules that we recently described in *C. deuterogattii* (13), since this number was much smaller than those found for proteins (5) and RNA (9) in cryptococcal vesicles. Of the 13 previously described small molecule components of cryptococcal EVs, 2 were below the detection level (Val-Leu-Pro-Val-Pro and asperphenamate) in this study, so we concentrated our analysis on the remaining 11 compounds. To compare the small molecule composition of EVs from WT and mutant cells, we used the <u>p</u>artial least <u>s</u>quare <u>d</u>iscriminant <u>a</u>nalysis (PLS-DA). This analysis allows the comparison of data including several variables under an algorithm-supervised mode, as is the case of mass spectrometry (MS) data. Different chemical profiles of small molecule composition of EVs from WT and mutant cells were found (Figure 7). As anticipated, the composition of the EVs from the *nop1*6Δ.1 and *nop1*6Δ.2 mutant strains was similar. However, EV composition in WT cells was clearly different from that observed in mutant vesicles. Three of the annotated metabolites were classified as variables important in the projection (VIPs) identified by PLS-DA, namely Phe-Pro, Pyro Glu-Leu, and Cyclo (Tyr-Pro). They were used for the discrimination between the three groups (small molecule composition of EVs produced by WT, *nop1*6Δ.1, and *nop1*6Δ.2 cells). These results proved the concept that deletion of Nop16 affected EV formation in *C. deuterogattii*.

**Figure 7.**
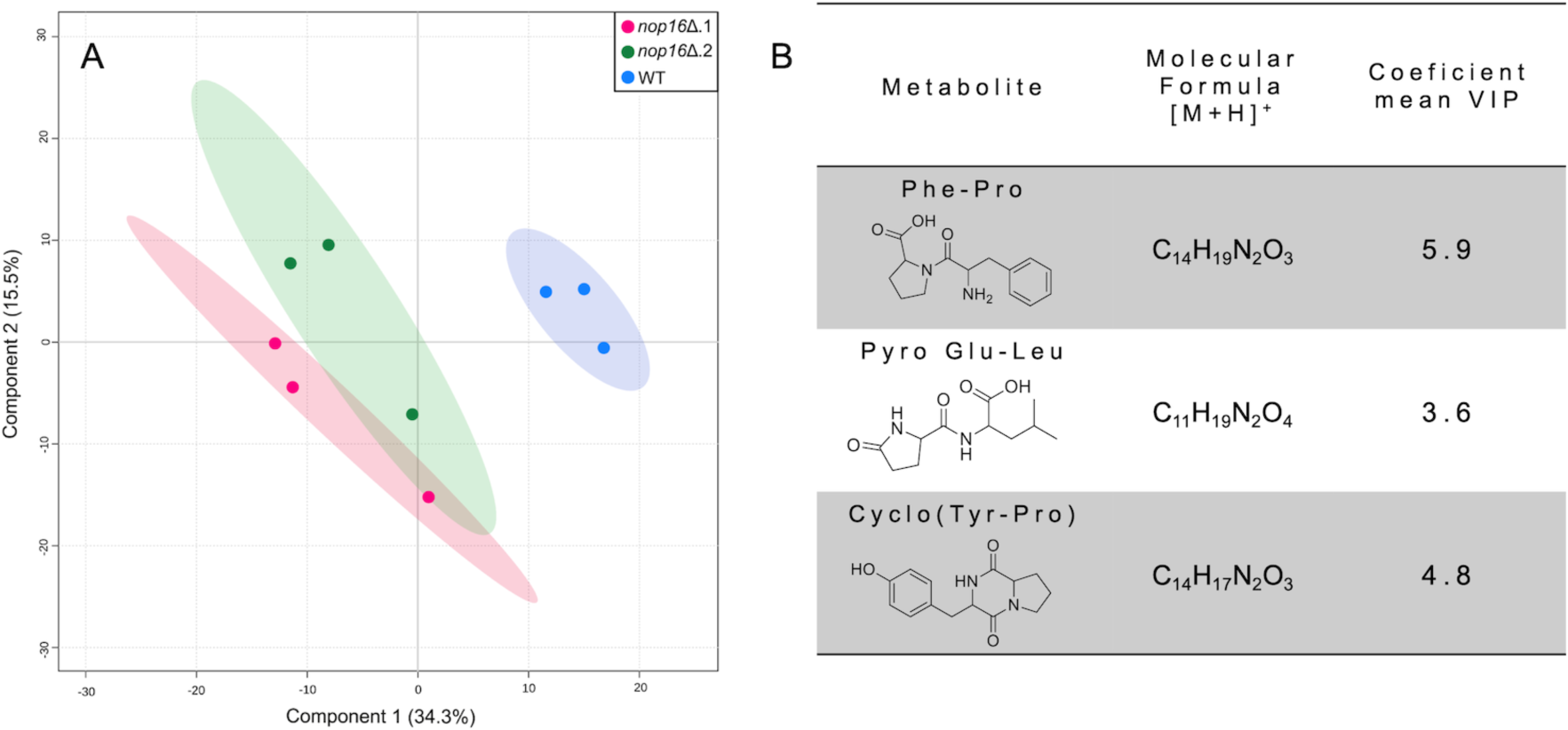
Multivariate data analysis of small molecule composition in WT and mutant cells of *C. deuterogattii* lacking expression of NOP16 (*nop1*6Δ.1 and *nop1*6Δ.2 strains) based on LC-MS data. A. <u>P</u>artial least <u>s</u>quare <u>d</u>iscriminant <u>a</u>nalysis (PLS-DA) of EVs LC-MS data. Each sphere in this analysis represents a single replicate. The elliptical area around the three replicates represents the confident zone of each group. In this analysis, R^2^ and Q^2^ values corresponded to 0.98 and 0.20, respectively, confirming the consistency between the original and cross-validation predicted data. B. Three metabolites [Phe-Pro, Pyro Glu-Leu, and Cyclo (Tyr-Pro)] were classified as <u>v</u>ariables important in the <u>p</u>rojection (VIP) features indicated by PLS-DA. Only VIPs with a coefficient score above 3.0 and p < 0.01 for the permutation test were selected. Their molecular formula [M+H]^+^ and coefficient scores calculated for the PLS-DA model shown in (A) are listed.

### EVs from parental cells increase the virulence of the *nop1*6Δ mutant cells

Since the EVs were altered under conditions of interrupted Nop16 expression, we asked whether the vesicles were the elements required for virulence in the *nop1*6Δ.1 and *nop1*6Δ.2 mutant strains. We raised two hypotheses. First, the hypovirulent profile of the mutants could be a result of the reduced number of EVs. Second, the altered composition of EVs would be responsible for the hypovirulent phenotypes. To address these hypotheses, we returned to the *G. mellonella* model of infection using the *nop1*6Δ.2 mutant, in which the formation of EVs was more evidently reduced. To test whether reduced EV concentrations would be the reason for the hypovirulence, we infected *G. mellonella* with *nop1*6Δ.2 cells supplementing the injecting suspension with EVs produced by this mutant strain. The amount of EVs added to this suspension was calculated on the basis that this mutant was 3-fold less effective in producing EVs than WT cells. Therefore, for each cryptococcal cell used for infection we added 3 EVs produced by the mutant. Independently on the presence of the EVs produced by the mutant strain, all larvae died by day 8 post-infection, and no significant differences were observed between systems injected with EVs plus the mutant or those receiving the mutant strain only (Figure 8, P = 0.5488). We then speculated that the number of EVs was likely not determinant for pathogenesis. To test whether the composition of EVs influenced virulence, we injected *G. mellonella* with the *nop1*6Δ.2 strain in the presence of EVs produced by WT cells, using the same ratio of 3 EVs for each cryptococcal cell used for infection. In comparison with systems receiving no EVs or EVs from mutant cells, a clearly accelerated death curve was observed when *G. mellonella* was injected with the *nop1*6Δ.2 mutant in the presence of EVs produced by WT cells (Figure 8, P = 0.0007).

**Figure 8.**
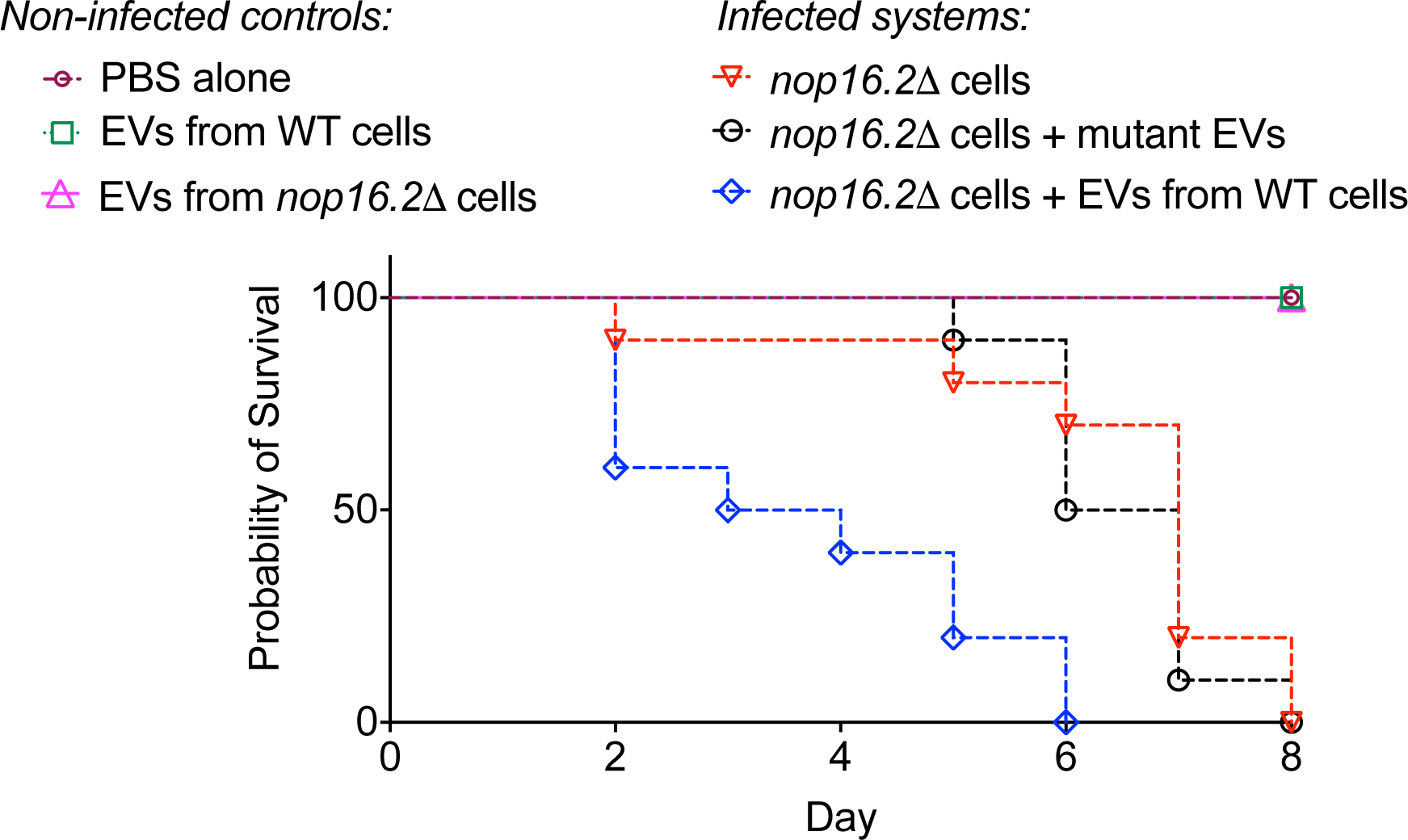
A role for EVs in the ability of *nop1*6Δ.2 mutant cells to kill *G. mellonella*. All larvae survived after injection with PBS alone, EVs from wild-type (WT) cells, or EVs from the *nop1*6Δ.2 mutant. Animals infected with the *nop1*6Δ.2 mutant alone or the mutant in the presence of their own EVs had similar mortality rates (P = 0.5488). In contrast, infection of *G. mellonella* with the *nop1*6Δ.2 mutant in the presence of EVs produced by WT cells resulted in higher mortality rates in comparison with all other systems infected with the *nop1*6Δ.2 mutant of *C. deuterogattii* (P = 0.0007). Statistical analysis was performed with the Mantel-Cox test.

## Discussion

In this study, our results connected the *NOP16* gene with EV biogenesis in *C. deuterogattii*, as concluded from the observation that the mutant strains with defective Nop16 expression produced fewer vesicles, and their small molecule composition was altered in comparison with wild-type cells. A role for a nucleolar protein in EV biogenesis might sound illogical at first sight. However, ribosomal proteins are major components of fungal EVs (5, 20). According to the recently launched ExVe tool for analysis of fungal EVs (34), at least 82 ribosomal proteins are present in *C. deuterogattii* EVs, with 17 of them being directly associated with the 60S ribosomal subunit. These results might have a relationship with the observation that, in *S. cerevisiae*, Nop16 was required for the biogenesis of the 60S ribosomal subunit (18, 19). If Nop16 plays a similar role in *C. deuterogattii*, its defective expression is expected to induce an altered EV cargo. To prove this concept, we analyzed the small molecule composition of *C. deuterogattii* EVs and confirmed that *NOP16* deletion affected the vesicular cargo. Other components, however, were not analyzed.

Our results indicated a connection between regular EV production and cryptococcal virulence. In this sense, it is noteworthy that the current literature indicates that EVs play puzzling roles in cryptococcal pathogenesis. These structures were denominated ‘virulence bags’ due to their complex composition that includes several virulence factors (20), which agrees with studies suggesting that cryptococcal EVs function in favor of disease progress. For instance, *C. neoformans* EVs facilitated cryptococcal traversal across the blood-brain barrier in mice, promoting an enhanced infection of the brain (36). An RNAi mutant strain of *C. neoformans* lacking expression of Sec6, a gene regulating conventional secretion, had attenuated virulence in mice in association with the absence of EV formation (37). More recently, long-distance communication via EVs was demonstrated to be essential for the virulence of *C. deuterogattii* (6).

Paradoxically, several studies indicate that cryptococcal EVs stimulate disease control. EVs stimulated the antifungal activity of macrophages in association with nitric oxide and cytokine production (4). In addition, EVs enriched with sterylglucosides induced protection against *C. neoformans* in a *G. mellonella* model of infection (31). Similarly, an EV peptide produced by *C. deuterogattii* induced the control of animal mortality in the same model (13). In mice, exposure to EVs produced by an acapsular strain of *C. neoformans* resulted in attenuated disease (5), which agrees with reports in the *C. albicans* model (39). The versatility of EVs in cryptococcal pathogenesis was efficiently illustrated in a recent study that demonstrated that vesicle properties changed according to the growth condition (40). EVs produced under a poor nutritional condition contained more virulence compounds, and induced a more robust inflammatory pattern than those produced in a rich nutritional medium. On the other hand, EVs produced in a rich medium inhibited the expression of genes related to the inflammasome, suggesting an involvement of EVs in the pathogenic process (40). We demonstrated recently that cryptococcal EVs are highly diverse in their physical-chemical properties, suggesting that their functions can indeed show a high diversity (33).

Our results suggest that an altered composition of EVs can result in attenuated virulence, as concluded from the observation that EVs produced by wild-type cells restored, at least partially, the virulence of a mutant strain lacking expression of Nop16. On the basis of the previous reports demonstrating that EVs contribute to the pathogenic potential of *Cryptococcus* (6, 20, 36, 37, 40), we speculate that the hypovirulent phenotype of the *nop16!1* mutants could be a result of altered EV production. Our results suggested that increasing the concentration of EVs was not sufficient to restore the pathogenic potential of the *nop16!1* mutants, since the mortality of *G. mellonella* did not change in the presence of an increased amount of EVs produced by these cells. In contrast, infection with the mutant cells in the presence of EVs produced by the parental cells resulted in an increased pathogenic potential. These results strongly suggest that EV cargo is determinant for the ability of *C. deuterogattii* to kill *G. mellonella*. One limitation of our study comes from the lack of knowledge of the EV components impacting virulence, since it is very likely that the composition of the vesicles produced by the *nop16!1* mutants was altered at multiple levels, and not only in the small molecule cargo. Regardless, these results might represent a proof of concept that EV formation is connected to virulence through their composition. Future studies identifying bioactive EV components and determining their vesicular concentration will be necessary for the characterization of the molecules impacting infection, as recently described for the Ile-Pro-Ile tripeptide (13). However, we anticipate that this might be a complex analysis since we cannot anticipate whether a molecular association between different vesicle components will be required for fungal virulence.

## Methods

### Strains

Fungal strains were maintained on Sabouraud agar plates (1% yeast extract, 2% peptone, 4% dextrose, and 1.5% agar) grown at 30°C for 24 h and stored at 4°C. Twenty-four h before the experiments, the strains were transferred to liquid YPD medium (1% yeast extract, 2% peptone, and 2% dextrose) and incubated at 30°C for 24 h with shaking (200 rpm). In experiments where capsule induction was necessary, the cells underwent an extra step of incubation in the capsule induction medium (Sabouraud 10% diluted in MOPS 50 mM, pH 7.4) (41) for 24 h at 37°C and 5% CO_2_. For the GXM detection assay by ELISA, cells grown in YPD were submitted to an incubation step in RPMI (37°C, 5% CO_2_ for 24 h) before the collection of the supernatants. The Delsgate methodology was used to construct the *NOP16* gene inactivation allele by employing the vector pDONR-NAT, as described previously (22). Primers used to amplify the *NOP16* gene flanking regions, as well as the internal fragment, are listed in Table 1. For the confirmation of *NOP16* deletion, the strains were cultured in YPD for 24 h with shaking (200 rpm). The cells were washed with fresh YPD and counted in a hemacytometer. A total of 500.000 cells were inoculated into fresh YPD in the absence (control) or presence of mebendazole (1 mM) in a 96-wells plate. After incubation for 24 h at 30°C, the OD600 was determined and the relative growth was determined by the ratio of optical densities in the presence of mebendazole by the control condition. For determination of growth rates, WT and mutant strains were grown in Sabouraud broth overnight at 30°C with shaking (200 rpm). The cultures were washed 3 times with PBS and counted in a Neubauer chamber. The cells were suspended (2.5 x 10^5^ cells/mL) in liquid Sabouraud, YPD, or Roswell Park Memorial Institute (RPMI) medium supplemented with 1% glucose, buffered (pH 7.0), with 165 mM morpholinepropanesulfonic acid (MOPS). Each suspension (200 μL, triplicates) was placed onto the wells of 96-wells plates and incubated at 37° C on a Molecular Devices SpectraMax Paradigm microplate reader. Fungal growth was determined with optical density measurements at 530 nm every hour for 48 hours.

**Table 1:**
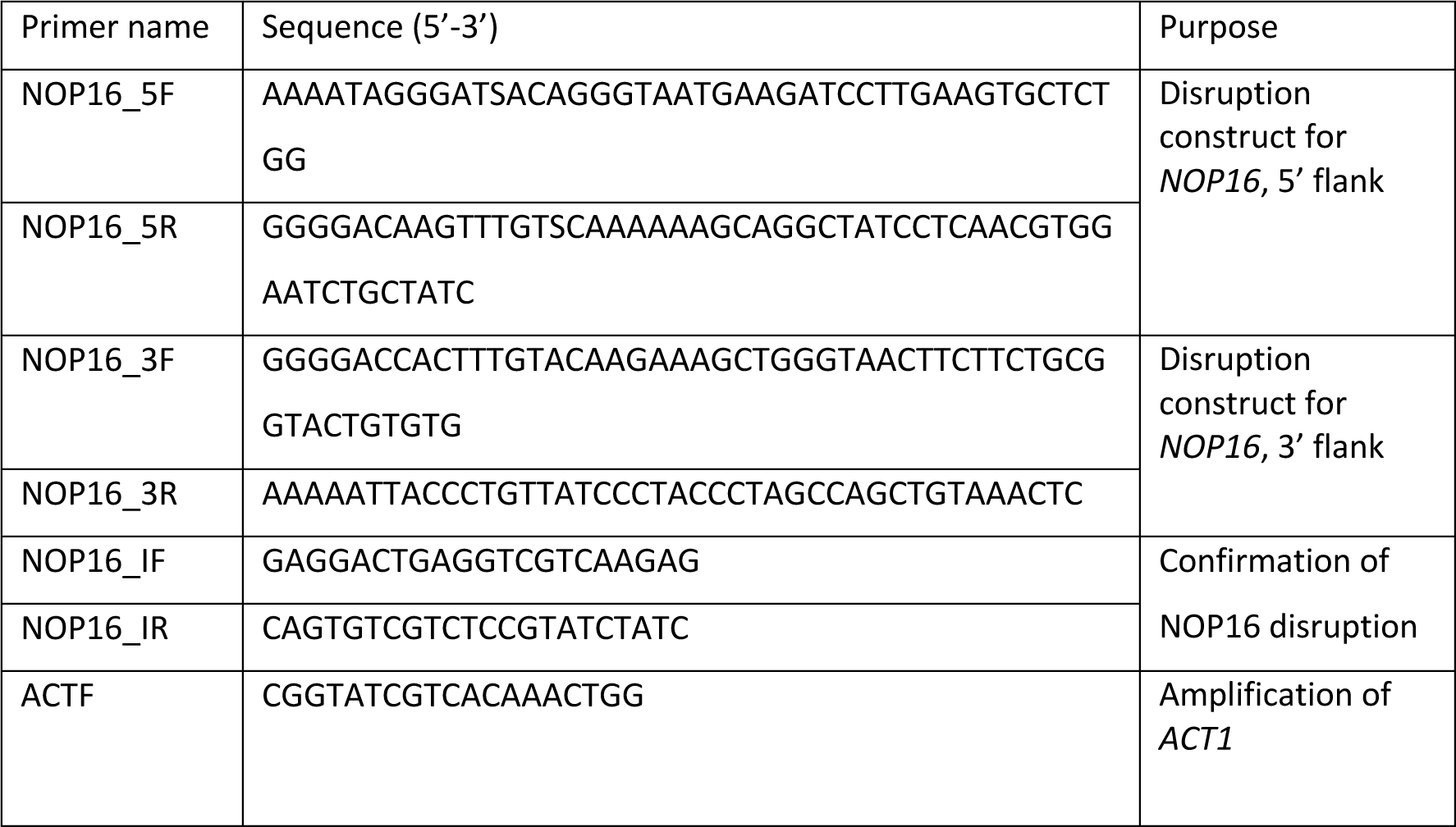
Primers used in this study

### Ultrastructure of *C. deuterogattii*

For the analysis of possible cellular alterations resulting from *NOP16* deletion, fungal cells were processed for transmission electron microscopy (TEM) (32). The cells were fixed in 4% paraformaldehyde and 2.5% glutaraldehyde in 0.1 M cacodylate buffer, pH 7.2 for 1 h at room temperature. The fixed samples were washed 3 times with cacodylate buffer by centrifugation (5,000 x g for 2 minutes). The pellet was treated with 1% osmium tetroxide and 5 mM potassium ferrocyanide diluted in 0.1 M cacodylate buffer (1:1 ratio) for 1 h at room temperature. The samples were then washed 3 times with cacodylate buffer and then sequentially dehydrated with 30%, 50%, 70%, 90%, and twice with 100% acetone. In each step, the samples were incubated with acetone for 30 minutes. The cell pellet was covered with the EMbed 812 resin (EMS) diluted in acetone. The initial dilution was 1 part resin to 2 parts 100% acetone, and this proportion was changed daily for three days (1 part resin to 1 part acetone on the second day; 2 parts resin to 1 part acetone on the third day). On the fourth day, the solution was replaced with 100% resin. This process was repeated, and the cell pellet covered with resin was incubated at 60°C for 48 h until complete polymerization of the resin was achieved. Ultra-sections were prepared with an ultramicrotome (Leica EM UC6; thickness of 70-80 nm) and mounted on microscopic grids. The observation of the samples was performed using a JEOL 1400Plus microscope with an acceleration of 90 kV. The images were obtained using a digital camera with an 8-megapixel CCD coupled to the equipment.

### Analysis of the polysaccharide capsule

Fungal cells were first counterstained with India ink for visualization of the capsule by light microscopy. As the polysaccharide structure is not permeable to India ink, counterstaining generates a high-contrast zone that allows the capsular structure to be visualized. *C. deuterogattii* cells were incubated in 10% Sabouraud in 50 mM morpholinopropanesulfonic acid (MOPS), pH 7.4 (41) for 24 h at 37°C and 5% CO_2_. After capsule induction, the cells were recovered by centrifugation at 3,000 x *g*, washed with PBS, and fixed with 4% paraformaldehyde. The cell suspension (3 µl) was placed onto a glass slide and supplemented with 2 µl of Indian ink. The sample was finally covered with a glass coverslip and observed with an inverted microscope (Leica DMi8). The images obtained in this process were analyzed using the ImageJ software (NIH). The capsule size was estimated as the cell body diameter subtracted from the total capsule diameter. For serological detection of capsular GXM, cryptococcal cells grown in YPD for 24 h were centrifuged and the cell pellet washed with PBS. The cells were transferred to the capsule induction medium under the conditions described above. The cells were then centrifuged at 3,000 x *g* for 3 minutes and the pellet was washed 3 times with sterile PBS. The cells were suspended in 150 µl of 4% paraformaldehyde in PBS and incubated at room temperature for 30 minutes. After fixation, the cell suspension was centrifuged again at 3,000 x g for 3 minutes. The cell pellet was subjected to three washes with PBS and blocked with 1 ml of blocking buffer (1% BSA in PBS), followed by incubation for 1 h at 37°C. The suspension was centrifuged, the supernatant was discarded and 150 µl of 25 µM calcofluor was added to the cell pellet, followed by incubation for 30 minutes at 37°C. Then, 3 washes were performed with PBS and 120 µl of an anti-GXM antibody 18B7 (donated by Dr. Arturo Casadevall, Johns Hopkins University, Baltimore, USA) at 10 µg/ml in blocking buffer were added to the cell suspension, for subsequent incubation for 1 h at 37°C. The cells were washed 3 times with PBS and suspended in a solution of 120 µl of a secondary antibody (goat to mouse immunoglobulin antibody) conjugated to Alexa 546, diluted in blocking buffer (1:1,500). The cells were incubated for 1 h in the dark at room temperature. The cells were washed again with PBS and stained with fluorescein-isothiocyanate (FITC) labeled wheat germ agglutinin (WGA) for detection of chitin oligomers in the cell wall (24). This step was carried out by suspending the cell pellet in 600 μl of WGA at 5 μg/ml, followed by incubation for 30 minutes at 37°C. The cells were washed with PBS and observed with an inverted fluorescence microscope DMi8 (Leica Microsystems). Extracellular GXM was measured by ELISA as previously described (42) with minor modifications (3).

The surface of fungal cells was also analyzed by scanning electron microscopy (SEM) (32). The *C. deuterogattii* cells grown in YPD were centrifuged at 3,000 x *g* and the cell pellet was washed with PBS. The sample was fixed with 2.5% glutaraldehyde in cacodylate buffer (0.1 M sodium cacodylate, pH 7.2). The cells were washed 3 times with a post fixative solution (0.2 M sucrose, 0.1 M sodium cacodylate buffer, 2 mM MgCl2). The washed samples (150 µl) were allowed to adhere to 0.01% poly-L-lysine (type I)-coated coverslips for 30 minutes at room temperature. The coverslips were sequentially dehydrated with 30%, 50%, and 70% ethanol solutions (5 minutes for each concentration), followed by 90% ethanol and two rounds of 100% ethanol, with these three steps lasting 10 minutes each. The cells were dried on a critical point chamber (Leica EM CPD300) and coated with gold particles on a metallizer (Leica EM ACE200). The visualization of the samples was performed using a JEOL JSM-6010 Plus/LA microscope with an acceleration of 5 kV.

### Golgi staining

We followed the protocol originally described by Pagano and colleagues (29) and adapted by our group to the analysis of the Golgi in cryptococci (30, 31). Briefly, yeast cells (10^7^) were fixed with 4% paraformaldehyde in PBS, followed by washing with PBS and incubation with C6-NBD-ceramide (10 µM) for 16 h at 4°C. The cells were then incubated with fetal calf serum (10%) at 4°C for 1h to remove the excess of C6-NBD-ceramide. The cell wall was stained with calcofluor white (0.1 mg/ml) for 30 minutes at room temperature, followed by washing with PBS and analysis by fluorescence microscopy as previously described (30, 31). Virtually all cells were efficiently stained with calcofluor white, and the percentage of cells giving positive signals for C6-NBD-ceramide-derived fluorescence was manually determined in populations of 100 cells for each tested condition.

### Isolation of EVs

The isolation of EVs produced by mutant and WT *C. deuterogattii* cells followed our recently described protocol (32). Fungal cultures had their cell densities determined in a Neubauer chamber and adjusted to 3.5 x 10^7^ cells/ml. Aliquots of each inoculum (300 µl) were spread with a Drigalski loop onto Petri dishes containing 25 ml of solid YPD. Three plates per strain were used. The plates were incubated at 30°C for 24 h to reach confluence, and then the fungal cells were recovered by gently scraping with an inoculation loop, and transferred to a centrifuge tube containing 30 ml of sterile PBS. Sequential centrifugation was used to remove the cells. In the first centrifugation step, the samples were centrifuged at 5,000 x g for 15 minutes at 4°C. The supernatants from the first centrifugation step were transferred to sterile tubes. These tubes were submitted to a second centrifugation step at 15,000 x g for 15 minutes at 4°C, for sedimentation of the cellular debris. After this step, the supernatants were filtered through 0.45 μm membranes and ultracentrifuged at 100,000 x g for 1 h at 4°C. The pelleted EVs were suspended in a final volume of 300 µl of PBS and stored at 4°C.

### Quantification and morphological analysis of EVs

The quantification of EVs was performed by nanoparticle tracking analysis (NTA) using LM10 nanoparticle analysis system coupled to a 488 nm laser, equipped with a camera and flow pump (Malvern Panalytical, Malvern, United Kingdom) and the NTA 3.0 software (Malvern Panalytical) (32). The samples were diluted 200-fold in PBS to achieve the optimal readout range of 9 x 10^7^ to 2.9 x 10^9^ particles/ml). The samples were injected with 1 ml syringes attached to a continuous flow injection pump. Three 60-second videos (camera level at 15, gain at 3) were obtained per sample after the passage of the samples through the light beam. The viscosity of the samples was indicated as the same as that of water. For data analysis, the camera gain was changed to 10-15 and the detection limit used was 3 for all samples. Transmission electron microscopy (TEM) was used to visualize the EVs (32). The samples were homogenized with a vortex for two minutes to break up possible aggregates. The EV suspensions (50 μl) were adhered to Formvar-coated grids for 60 minutes at room temperature. The grids were then washed with 30 µl of sterile PBS. The excess buffer was removed by applying filter paper to the bottom of each grid. The grids were then incubated with 30 μl of the Karnowski solution for 10 minutes, washed 3 times with cacodylate buffer, and finally dried with filter paper. The samples were counterstained with 5% uranyl acetate for 2 minutes. The grids were washed once with H2O, dried with filter paper, and transferred to a metallizer (Leica EM ACE200), where they were covered with carbon particles for later visualization with a JEOL 1400Plus microscope with beam acceleration at 90 kV.

### Urease activity

Cell suspensions at 1 x 10^8^ cells/mL were prepared in urea broth/Robert’s medium (final volume of 1 mL). The systems were incubated overnight at 30°C with shaking. The cells were pelleted by centrifugation and the supernatants (200 µl) were transferred to the wells of 96-well plates. Optical densities at 540 nm were determined on a SpectraMax PARADIGM microplate reader (Molecular Devices).

#### *C. deuterogattii* phagocytosis

Murine macrophage cells (RAW 264.7) were maintained in Dulbecco’s modified Eagle’s medium (DMEM) supplemented with 10% fetal bovine serum (FBS) at 37°C in 5% CO_2_. On the day before the experiment, a suspension containing 5 x 10^5^ cells/ml was prepared in DMEM supplemented with 10% FBS. 200 µl of this suspension were added to each well of 96-well plates. The plates were incubated at 37°C and 5% CO_2_ for 24 h. Fungal cells were adjusted to a density of 1 x 10^6^ cells/ml in YPD, washed once with PBS, and then incubated with 0.5 mg/ml FITC in PBS at room temperature for 15 minutes in the dark. The cells were washed 3 times with PBS and then the inoculum was adjusted to 5 x 10^5^ cells/ml in DMEM. The 18B7 anti-GXM antibody was added to a final concentration of 10 µg/ml, and opsonization was carried out by incubating these suspensions for 1 h at 37°C in a 5% CO_2_ atmosphere. Opsonized cells were used to infect macrophages previously seeded in 96-well plates, using the MOI (multiplicity of infection) of 1:1. The systems were incubated for 3 h at 37°C in 5% CO_2_. After incubation, the wells were washed 3 times with PBS to remove free fungal cells and 200 µl of DMEM + 10% FBS was added to each well. The plates were then transferred to an Operetta high-content imaging system (PerkinElmer) (43). The equipment was adjusted to a temperature of 37°C with a 5% CO_2_ atmosphere, and programmed to automatically photograph the infected cells. The images were obtained in the Alexa 488 channels to detect the fluorescence emitted by the FITC, and also under the bright field mode. The images were processed using the Harmony high-content imaging and analysis software (PerkinElmer) and ImageJ (NIH). Phagocytosis indices consisted of the percentage of cells infected with the fungus in each field.

### Galleria mellonella infection

To assess the pathogenic potential of the WT and mutant strains, the invertebrate model of infection in *Galleria mellonella*, which was validated for the study of cryptococcal pathogenesis (44), was used. For acclimatization, *G. mellonella* larvae weighing between 0.10-0.15 g were divided into groups of 15 animals in Petri dishes and incubated overnight at 37°C before infection. Suspensions containing 1 x 10^8^ cells/ml were prepared in sterile PBS for each of the strains to be analyzed. 10 µl of these suspensions, containing 1 x 10^6^ fungi, were used to infect the larvae of *G. mellonella* using a Hamilton syringe as previously described (13). The plates were then incubated at 37°C to monitor the survival of the animals over the days. The groups analyzed were: 1) animals infected with the WT, R265 strain; 2) animals infected with the *nop16*!1.1 mutant strain; 3) animals infected with the mutant strain *nop16*!1.2 and 4) control group inoculated only with PBS. Larvae mortality was observed daily, as evidenced by the lack of movement after stimulation with forceps.

*G. mellonella* mortality was also the model used to evaluate the role of EVs on fungal pathogenesis. In these assays, all infection conditions were similar to those described above. However, we only used the *nop1*6Δ.2 mutant, based on its lower ability to produce EVs. Six groups were analyzed, including non-infected controls (larvae injected with i) PBS alone, ii) EVs from WT cells, or iii) EVs from the *nop1*6Δ.2 mutant) and infected systems (infection with i) *nop1*6Δ.2 cells alone, ii) *nop1*6Δ.2 cells with EVs from WT cells, or iii) the *nop1*6Δ.2 mutant with its own EVs). The amount of externally supplemented EVs corresponded to 3 vesicles per each of the 1 x 10^6^ fungi used to infect *G. mellonella*. Monitoring of mortality was performed as described above.

### Small molecule analysis

*C. deuterogattii* EVs were prepared and analyzed as recently described by our group (13, 14). Briefly, the samples were vacuum dried, extracted with methanol, filtered through 0.22 μm membranes, dried under a N2 flux and stored at −20°C. The vesicular extracts were suspended in MeOH and submitted to ultra-high-performance liquid chromatography coupled to mass spectrometry (UHPLC-MS) on a Thermo Scientific QExactive^®^ hybrid Quadrupole-Orbitrap mass spectrometer. The parameters used for this analysis, including those used for tandem Mass spectrometry (MS/MS), were recently detailed by our group. UHPLC-MS operation and spectra analyses were performed using Xcalibur software (version 3.0.63). A molecular network was created using the online workflow (https://ccms-ucsd.github.io/GNPSDocumentation/) on the GNPS website (http://gnps.ucsd.edu) as reported in our recent study (13).

### LC-MS data analysis for multivariate data analysis

After Molecular Networking analysis on the GNPS database, the online visualization option was selected to process the original files. The LC-MS chromatograms were analyzed in the Feature finding option using MZMine2 (Dashboard online GNPS version). The following parameters were used: Precursor tolerance of 10 ppm; Noise level of 10E^4^; minimum and maximum peak width: 0.05 –1.5 min; and a retention time tolerance of 0.3 min. The job can be accessed online at: (https://gnps.ucsd.edu/ProteoSAFe/status.jsp?task=395c343397384433998e001434bedfed). The quantification table (.csv) was submitted to MetaboAnalyst 5.0 (https://www.metaboanalyst.ca/) for Statistical Analysis, and the data was processed using sum normalization and Pareto Scaling. Principal Component Analysis (PCA) and Partial Least Square-Discriminant Analysis (PLS-DA) were performed on the dataset. For the individual metabolite quantification, the interesting features (annotated *m/z*) were searched in the Xcalibur software (version 3.0.63), through Extracted Ion Chromatogram (EIC) analysis and integrated.

### Statistical analysis

Statistical analyzes were performed using the GraphPad Prism software 9.3 (GraphPad Software, Inc. La Jolla, USA). The results were analyzed using a one-way ANOVA and Tukey’s post hoc tests, except for the survival assay with *G. mellonella*, where the analysis with Mantel-Cox was used. The differences found were considered significant when the P values were smaller than 0.05.

## Acknowledgements

M.L.R. is supported by grants from the Brazilian Ministry of Health (grant 440015/2018-9), Conselho Nacional de Desenvolvimento Científico e Tecnológico (CNPq; grants 405520/2018-2 and 301304/2017-3), and Fiocruz (grants PROEP-ICC 442186/2019-3, VPPCB-007-FIO-18, and VPPIS-001-FIO18). M.L.R. also acknowledges support from the Instituto Nacional de Ciência e Tecnologia de Inovação em Doenças de Populações Negligenciadas (INCT-IDPN). H.C.O. received scholarships from the Inova Program of Fiocruz. F.C.G.R. received a scholarship from the Coordenação de Aperfeiçoamento de Pessoal de Nível Superior (CAPES, Brazil, Finance Code 001). T.P.F. acknowledges support from FAPESP, grant number 2021/00728-0. The funders had no role in the decision to publish, or preparation of the manuscript. The authors are grateful to the Program for Technological Development in Tools for Health-RPT-FIOCRUZ for using the microscopy facility, RPT07C, Carlos Chagas Institute, Fiocruz-Paraná. M.L.R. is currently on leave from the position of associate professor at the Microbiology Institute of the Federal University of Rio de Janeiro, Brazil.

